# AD Informer Set: Chemical tools to facilitate Alzheimer’s disease drug discovery

**DOI:** 10.1101/2021.07.22.453404

**Authors:** Frances M. Potjewyd, Joel K. Annor-Gyamfi, Jeffrey Aubé, Shaoyou Chu, Ivie L. Conlon, Kevin J. Frankowski, Shiva K. R. Guduru, Brian P. Hardy, Megan D. Hopkins, Chizuru Kinoshita, Dmitri B. Kireev, Emily R. Mason, Charles Travis Moerk, Felix Nwogbo, Kenneth H. Pearce, Timothy Richardson, David A. Rogers, Disha M. Soni, Michael Stashko, Xiaodong Wang, Carrow Wells, Timothy M. Willson, Stephen V. Frye, Jessica E. Young, Alison D. Axtman

**Author notes:** Corresponding author; Alison D. Axtman, UNC Eshelman School of Pharmacy, Division of Chemical Biology and Medicinal Chemistry, Structural Genomics Consortium, Chapel Hill, NC, USA.

## Abstract

**Introduction:** The portfolio of novel targets to treat Alzheimer’s disease (AD) has been enriched by the AMP-AD program.

**Methods:** A cheminformatics-driven effort enabled identification of existing small molecule modulators for many protein targets nominated by AMP-AD and suitable positive control compounds to be included in the set.

**Results:** We have built an annotated set of 171 small molecule modulators, including mostly inhibitors, targeting 98 unique proteins that have been nominated by AMP-AD consortium members as novel targets for AD treatment. These small molecules vary in their quality and should be considered chemical tools that can be used in efforts to validate therapeutic hypotheses, but which would require further optimization. A physical copy of the AD Informer Set can be ordered via the AD Knowledge Portal.

**Discussion:** Small molecule tools that enable target validation are important tools for the translation of novel hypotheses into viable therapeutic strategies for AD.

## 1. INTRODUCTION

Current approved treatments for Alzheimer’s disease (AD) address late-stage symptomatic effects of the disease [1, 2]. The discovery of disease-modifying interventions that slow or halt progression of AD are critical unmet needs. Promisingly, there has been a recent increase in the number of novel therapeutic strategies targeting AD that have progressed to clinical trials and programs have been developed to foster the discovery of novel targets implicated in AD pathology [3].

The National Institute on Aging (NIA), recognizing the need to explore new therapeutics for AD, established the Accelerating Medicines Partnership AD (AMP-AD) [4] precompetitive partnership among government, industry, and nonprofit organizations in 2014 to transform the existing model for AD diagnostic and drug development. At its inception, the goal of AMP-AD was the discovery of novel, clinically relevant therapeutic targets, and development of biomarkers to help validate current therapeutic targets [5]. AMP-AD was initiated as two arms: the Biomarkers in Clinical Trials Project and the Target Discovery and Preclinical Validation Project. The AMP-AD Biomarkers Project supported two Phase II/III secondary prevention trials testing anti-amyloid therapies with the goal of exploring the utility of tau imaging for tracking responsiveness to treatment and/or disease progression [5].

The AMP-AD Target Discovery and Preclinical Validation Project aimed to accelerate the AD drug discovery process via integration of analyses of large-scale molecular data from human brain samples with network modeling approaches and experimental validation [5]. This arm funded six multi-institutional, multidisciplinary teams who analyzed and integrated multidimensional human “omic” data (genomic, epigenomic, RNAseq, and proteomic) from more than 2000 human brains at all stages of AD and matched controls with clinical and pathological data [5]. A fundamental deliverable of this program was open sharing of all data via the AMP-AD Knowledge Portal and Agora [5]. The types of data generated included novel therapeutic targets for AD, a systems-level understanding of the networks within which these novel targets operate, and evaluation of their druggability in multiple model organisms [5]. A round of NIA-sponsored AMP-AD enhancer grants in 2017 augmented the AMP-AD data infrastructure and analytical capabilities and enabled generation of new human multi-omic data from brain, CSF, and blood samples, as well as single nucleus sequencing data from human and mouse brains. The NIA-funded TaRget Enablement to Accelerate Therapy Development for AD (TREAT-AD) consortium was subsequently started in 2019 to translate AD molecular signatures generated by the AMP-AD groups into new treatments [6]. TREAT-AD is made up of two multi-institutional research centers that subscribe to the same open dissemination of all data, research methodologies, and computational and experimental tools [6]. Building on the more than 500 new candidate AD targets nominated by the AMP-AD teams, the TREAT-AD groups will develop a series of new therapeutic hypotheses centered around a prioritized set of the novel proposed targets and, importantly, deliver a suite of target enabling tools including high-quality antibodies and chemical probes [6]. At the inception of the program, two versions of target enabling packages (TEPs) will be created: basic and full [7]. A basic TEP includes a knockout cell line, validated antibodies, and purified protein related to a protein target of interest. A full TEP delivers the basic TEP components plus a biochemical and/or biophysical assay, crystal structures, an AD-relevant knockdown cell line, and initial chemical matter.

High-quality small molecules are powerful tools that enable interrogation of biological pathways and the exploration of pharmacology in various biological contexts. Through optimization for potent and selective interaction with a specific target in cell-based assays, the role of a protein in various biological contexts can be studied. The term chemical probe has been defined as meeting certain specific criteria, including thresholds related to on-target potency, family-wide and sub-family selectivity, and cell-based activity [8, 9]. The stringency of requirements to be considered a chemical probe are such that relatively few have been nominated, since each probe demands extensive resources to be designed, optimized, and characterized. Those that have been developed, however, have been profoundly impactful for the research community, especially when released with an open sharing policy [10-15].

The huge investment of resource required to deliver a chemical probe has motivated the generation of chemogenomic and other compound sets. The public kinase inhibitor sets (PKIS/PKIS2) were two early examples of protein-class targeted sets comprised of published kinase inhibitors that were shared openly alongside all available annotation with the research community [16, 17]. These sets, while not comprised of chemical probes, have been shared with over 300 laboratories and have resulted in many new research findings, grants, and publications [18]. Furthermore, experience with these sets informed assembly of the kinase chemogenomic set (KCGS), which only includes kinase inhibitors for nearly half of the human kinome that show potent kinase inhibition and a narrow spectrum of activity when screened across a large panel of kinase biochemical assays [19]. Like PKIS/PKIS2, KCGS is made available with all associated annotation to all interested investigators. Sets of epigenetic modifiers have also been assembled into larger compound libraries that are openly distributed [9, 20]. In this same vein, Boehringer Ingelheim has set up the opnMe program to openly share molecules [21]. An overarching theme is that the wide-reaching impact of these small molecules and compound sets is realized due to open sharing, enabling interested investigators to easily request the sets, use them without restrictions, and publish their results.

The Cancer Target Discovery and Development Network within the National Cancer Institute (NCI) developed the idea of an “Informer Set.” More than 350 small molecules were assembled and profiled across many human cancer cell lines to reveal dependencies. This Informer Set was comprised of 35 FDA-approved drugs, 54 clinical candidates, and 266 additional tool compounds, which were selected to target distinct nodes in cancer cell circuitry and collectively modulate a broad array of cellular processes. The resultant dataset was deposited into a publicly accessible data portal known as the Cancer Therapeutics Response Portal (CTRP) to be used by other researchers, such as to make connections between the genetic and lineage features of cancer cell lines and small-molecule sensitivities [22]. The Informer Set, which is a living resource, has since been expanded to include ∼545 molecules and corresponding data deposited in CTRP as before. While physical copies of the compound set are not a shared resource, the data within CTRP is immensely useful to the cancer-focused research community and once again demonstrates the impact of open sharing [23].

## 2. METHODOLOGY

To assemble an AD Informer Set of small molecules, protein targets for AD were collected from AMP-AD and prioritized by the TREAT-AD teams, most of which can be found within the AD Knowledge Portal [5]. From the start of this collaborative effort between the TREAT-AD teams, priorities for data collection were agreed upon, duplication was avoided, and data gathering expertise was shared. Next, an exploratory cascade to find small molecules was established, which involved searching the literature and available databases including the Chemical Probes Portal and ICR Probe Miner [9, 24]. While there are exceptions, criteria for selection of most small molecules included on-target biochemical potency, preferably with activity at <1 μM, and reported cell activity at <10 μM. It is important to note that the compounds selected for inclusion do not all meet the quality criteria for chemical probes because this would have been overly restrictive given the less-explored nature of many nominated targets. These small molecules have been characterized as inhibitors, substrates, activators, agonists, or antagonists and should be treated as modulators of a biological target implicated in AD. Like the NCI Informer Set, these compounds are in various stages of development, ranging from only *in vitro* studies; to use in animals; to approved drugs. To supplement these more investigational small molecules, an additional set of small molecule modulators of advanced AD targets were added as positive controls. These clinical phase compounds were included as more extensively characterized examples that have been used to validate therapeutic hypotheses and for which phenotypic data has already been published. Reported AD-relevant phenotypes for the targets of these positive control compounds as well as for all other protein targets where they could be found are incorporated into the annotation for the AD Informer Set. Predicted and experimentally collected data for either the entire set or a subset of compounds is also included. Finally, all AD Informer Set compounds have been validated for quality by ^1^H NMR and LC–MS and registered with this data within the UNC Laboratory Information Management System (LIMS) within the Center for Integrative Chemical Biology and Drug Discovery (CICBDD).

## 3. RESULTS

The AD Informer Set is a physical set of high-quality compounds with associated data that can be used by the scientific community to interrogate AD-implicated biology. The set is comprised of 171 small molecules that target 98 unique proteins. Where possible, more than one compound preferably built upon differing chemotypes was selected as a modulator of each protein target. Unsurprisingly, protein targets that have been more widely studied yielded more available chemical tools, whereas those within the lesser studied proteome provided fewer choices. This resulted in 1–7 compounds being included for each protein target, with at least two for more than half (51) of the 98 protein targets in the set. A total of eleven positive control compounds that are currently in Phase 2/3 clinical trials or already approved drugs were selected for the AD Informer Set. Including these positive control compounds results in a distribution of chemical tools in terms of stage of development. Within the AD Informer Set, 36 compounds are approved drugs, 40 are in clinical trials (Phases 1–4), 68 have advanced into animal-based studies, and 27 have not been explored beyond cellular studies. For all 171 compounds in the set, we have performed and included in accompanying annotation files prediction of blood–brain barrier (BBB) permeability, experimentally determined kinetic solubility measurements, and single-concentration (10 μM) results in cell-based assays related to microglial viability and phagocytosis. Non-toxic compounds for which an interesting phenotype was observed in the phagocytosis assay were followed up in dose-response. A subset of nine selected compounds within the AD Informer Set were prioritized by the Emory/Sage/SGC TREAT-AD team for in depth study. For these compounds, GPCR profiling, human induced pluripotent stem cell (iPSC)-derived cellular assays, and mouse pharmacokinetic (PK) data were collected. These more extensive studies helped establish workflows and benchmarks for AD chemical probe development within the Emory/Sage/SGC TREAT-AD active target portfolio.

### Physical set specifics

The AD Informer Set will be provided to interested investigators as a single 384-well plate containing 1 μL of each compound as a 10 mM stock in DMSO. A plate map, included in the Supporting Information, will be provided when the plate is shipped. All information about ordering the set can be found on the AD Knowledge Portal.

### Data annotation

Annotation for the entire set is included as Supporting Information and also uploaded on the AD Knowledge Portal. This spreadsheet is alphabetized by gene target and has been divided into two sections: compound-specific information and gene-specific information. Within the compound-specific information section, there is compound identification data including the assigned UNC number, associated lot ID, an alternative (common) name, CAS number, vendor that provided the solid sample, SMILES string, and designation as a positive control (where applicable). Also within this subsection is data related to predicted BBB penetration by the StarDrop software program (log([brain]:[blood])) [25-28] as well as by the ADMET Predictor software program [29-31]: qualitative likelihood of crossing BBB and logarithm of the brain/blood partition coefficient (LogBB). Other predicted compound properties included from the ADMET Predictor software program are whether the compound is a likely P-glycoprotein (P-gp) and/or breast cancer resistance protein (BCRP) substrate and/or inhibitor, making it susceptible to efflux. Also included is literature-reported data for the compounds, including on-target biochemical potency (in nM), whether the compound demonstrates cell activity when dosed at <10 μM, stage of development and dosing information, kinetic solubility (in μM), and single-concentration microglial viability and phagocytosis assay activity. For a subset of nine compounds, there is also an indication within this spreadsheet that it was included in GPCR profiling, iPSC-based and mouse PK studies. PubMed IDs and/or the appropriate database have been included as references for the potency and dosing information assembled.

Within the gene-specific information portion of this spreadsheet is the Ensembl gene ID(s), published AD-relevant phenotypes associated with gene, predicted AD therapeutic direction, and nomination details. Most genes were nominated by AMP-AD investigators, while others have been prioritized by TREAT-AD team members including those selected as positive controls. PubMed IDs and/or the appropriate database have been included as references for AD-relevant phenotypes referenced within the file. Several of the details related to the gene targets within this spreadsheet can be cross-referenced on the Agora website, which can be searched by Ensembl ID or gene symbol. This invaluable resource can be used to find additional specifics surrounding target nomination, evidence of association with AD, expression profiles within the brain, and predicted druggability.

### Kinetic solubility determination

Analyses were executed using 10 mM DMSO stocks of AD Informer Set compounds in aqueous buffer at neutral pH (7.4) by Analiza, Inc via total Chemiluminescent Nitrogen Determination (CLND) as previously described [32]. Since this method relies on nitrogen detection for quantification, no kinetic solubility value could be determined for compounds lacking nitrogen in their structure (16 in total). All calculated solubility values, after correction, are included in the annotation file in the Supporting Information. These values are provided to users of the set to make them aware of compounds with sub-optimal solubility that may result in precipitation in assays executed in aqueous buffers or media.

### Microglial viability and phagocytosis studies

All AD Informer Set compounds were tested in two immortalized microglial cell lines, mouse (BV2) and human (HMC3), in a high content imaging assay in duplicate by the Chu lab. Assays were carried out at a single concentration (10 μM), with plated cells allowed to adhere overnight, then treated the next morning (day 2), and pHrodo-myelin (labeled phagocytosis ligand, purified from mouse brain) seeding ∼8h later. Plates were then imaged on day 3, approximately 22 hours after pHrodo-myelin addition. Cytotoxicity was evaluated by cell counts per well, nuclear area, and nuclear DNA intensity. Phagocytosis was measured by mean total intensity of phagocytotic vesicles per cell. Idelalisib (10 μM) and cytochalasin-D (4 μM) were used as positive controls in the assay. All data, which was an average of 2 data points, was normalized to untreated wells and reported as % control, which is included in the Supporting Information both as a raw data file and summarized in the compound/gene-annotation file. A few compounds were observed to have inherent fluorescence, and these are noted in the Supporting Information files. Compounds that were not cytotoxic at 10 μM and showed a phenotype (inhibition or activation) were followed-up in dose-response. Based on the single-concentration data, several compounds were selected for 10-point dose-response follow-up in duplicate or triplicate. These compounds were initially dosed up to 20 μM for 24h and then the highest dose was adjusted to 40 μM, once again treating for 24h. Finally, a subset of compounds was dosed up to 40 μM for 48h. Single-point data was compared with the dose-response data.

### GPCR panel

A subset of nine selected compounds were submitted to the NIMH-sponsored Psychoactive Drug Screening Program (PDSP) at UNC. Screening against the receptors within the PDSP panel offers an assessment of off-target pharmacological activity at cloned human or rodent CNS receptors, channels, and transporters that could confound CNS-related phenotypes. A total of 46–47 primary binding assays were run in quadruplicate on these compounds at 10 μM, with follow-up secondary binding assays in 11-point dose-response in triplicate for select compounds versus select receptors [33]. Secondary binding assays were executed for all receptors where test compound displaced an average of >50% of radio-labeled standard ligand. As an exception, no secondary assays were carried out for H3, PBR, or GABAA. This data is included as Supporting Information.

### Human iPSC-derived cellular assays

hiPSC-derived neural cells are increasingly being used as preclinical models in neurodegenerative research [34]. With this concept in mind, the same nine compounds were preliminarily analyzed by the Young lab in a panel of AD-relevant assays. iPSC-derived neurons were differentiated and plated for assays as we have previously described [35, 36]. Neurons were treated with either 0.1 or 1 μM of compound for 24h or 48h, and media, cell lysates, and RNA were harvested. Basal secretion of Aβ peptides (Aβ40 and Aβ42) by iPSC-derived neurons into culture media was assessed at two time points (24h and 48h) using a multiplexed ELISA assay (Figure 3). We calculated the fold-change from the vehicle control (DMSO) measured for Aβ40 and Aβ42. To look at the relationship of these two peptides to each other, we calculated the Aβ42:40 ratio for each compound. Measurements of phospho-Tau(Thr231) and total Tau peptides were measured from neuronal lysates, also using a multiplexed ELISA assay. Using this assay, we measured the phospho:total tau ratio for each condition compared to vehicle (DMSO) treatment as well as the individual changes in phospho and total Tau peptides.

### PK data

The same nine compounds were also selected for mouse PK studies at Pharmaron. Since mouse PK data had previously been generated and published for three of these compounds (UNC2025C, UNC10302676A/ ABT-957 (alicapistat), and UNC10302675A/ MDL 28170) [37-39], the remaining six compounds were sent for snapshot PK. Pooled plasma was analyzed at 0.5h, 1h, 2h, and 4h following a single IV dose (3 mg/kg) in CD1 mice (2 mice per cohort) for each compound (Table 2). In parallel, the three compounds with known PK data were dosed IV at 10 mg/kg and plasma plus brain concentrations measured at 1h in CD1 mice (3 mice per cohort, Table 2). Published IV PK results for UNC10302676A/ ABT-957 (Alicapistat), UNC10302675A/ MDL 28170, and UNC2025C are also included in Table 2 for reference [37-39]. Based on PK snapshot results for the six compounds analyzed, three (UNC10302679A/ GSK2256294A, UNC10302681A/ TPPU, and UNC10302682A/ P505-15 (PRT06207 HCl)) were sent for brain exposure measurement as described above. UNC10302683A/ AR9281 was not checked for brain permeability due to its short half-life, while the solubility of UNC10302678A/ entospletinib (GS-9973) and UNC10302680A/ iMDK, requiring a formulation in NMP/PEG-400 (10:90) to execute the snapshot PK studies, excluded them in the analysis of brain permeability.

## 4. DISCUSSION

The intended uses of the AD Informer Set include: (1) to interrogate target validity in established and emerging AD models, (2) to serve as positive controls/comparators versus new chemical entities (NCEs) for AD, and (3) to qualify newly developed or established AD-relevant assays. Neuroinflammation, for example, is a phenotype that is increased early in AD and, because of its chronic nature, is considered pathogenic [40, 41]. Based on its known role in propagating AD pathology, neuroinflammation is a phenotype that could be investigated using the AD Informer Set. This set is not comprised of highly optimized, potent, and selective small molecule chemical probes [8, 9], and results generated when using the set should be treated accordingly. In particular, selectivity assessment is limited for many compounds and pharmacological results should be cross-validated wherever possible by genetic methods of target manipulation [42]. Most compounds contained herein were optimized within certain biological contexts for applications other than the treatment of AD. Many remain unoptimized. Thus, the set should be used to explore related pharmacology within the context of AD. AD Informer Set compounds could then serve as chemical starting points in medicinal chemistry campaigns focused on delivering AD-specific tools. The inclusion of 11 positive control compounds, which are in Phase 2/3 clinical trials or FDA-approved drugs developed for AD, however, allows benchmarking in specific AD phenotypic assays related to the pathways for which these advanced compounds were developed (see Supporting Information).

As acknowledged above, we have not acquired target class selectivity for the compounds included in the AD Informer Set, and they likely lack the requisite selectivity embodied by chemical probes. As AD is a complex and multifactorial disease and current therapies available show limited ability to modify the disease, the approach of multi-target drug design is being pursued to simultaneously modulate multiple targets [43]. Thus, the polypharmacology likely elicited by AD Informer Set compounds in phenotypic assays may reveal phenotypes worthy of further investigation.

Other attractive features of the AD Informer Set include the ability to execute an unbiased screen of compounds that modulate targets implicated in AD. As proof-of-concept and to demonstrate its utility, we have analyzed compounds within the larger set in cell-based assays involving immortalized mouse and human cells as well as human iPSC-derived neurons. Some unanticipated results were obtained when these assays were carried out, leading to the design of follow-up studies and new hypotheses about the gene targets as discussed below. Since the set is comprised of modulators that have differential effects on their target(s), including inhibition or activation, it is anticipated that some will transiently phenocopy AD pathology while others may offer a new therapeutic direction to pursue.

Beyond simply highlighting those compounds that will likely precipitate in aqueous buffers, the kinetic solubility study results provide general insights. As nearly all compounds (101 of 155) demonstrated solubility >10 μM, most compounds in the AD Informer Set are not considered poorly soluble. For reference, during a medicinal chemistry optimization program a kinetic solubility concentration >20 μM is often considered acceptable (93 of 155 compounds fit this criteria), while a concentration >100 μM is desirable (67 of 155 compounds fit this criteria). While drugs with low water solubility are predisposed to low and variable oral bioavailability and there are structural modifications known to improve aqueous solubility, ∼40% of currently marketed compounds remain poorly water-soluble [44-46]. As further support that approved drugs do not have to fit a certain kinetic solubility concentration, six of 36 approved drugs in the AD Informer Set have kinetic solubility values <10 μM and 15 of 36 have kinetic solubility values <100 μM. It has been suggested that kinetic solubility may also play a minor role in passive diffusion across the BBB [47].

Microglial viability studies demonstrated that most compounds in the set are not toxic. We chose to profile the AD Informer Set in both human (HMC3) and murine (BV2) immortalized microglia. While most compounds in the set were likely optimized in a human context, available *in vivo* model systems for AD are non-human. We expect that there will be target differences between humans and these model systems. Our use of both HMC3 and BV2 cell lines highlights that there is cell line to cell line as well as species to species differences in target expression and pathway connectivity.

The role of microglia in AD has been well documented and was thus chosen as a cell type in which to profile the AD Informer Set [48-50]. As an example, neuroinflammation, as detected by the presence of activated complement proteins, interleukins, cytokines, and chemokines in microglia and astrocytes, is increased early in AD and is considered pathogenic due to its chronic nature [40, 41]. Cytotoxic effects can be assessed by considering multiple parameters. A cell count <70% control (lower number, more toxic) is the simplest measurement. More nuanced details come from considering nuclear area measurement (<80% control indicates toxicity) and nuclear DNA intensity, where >120% control is indicative of early apoptosis and <70% is indicative of later apoptosis. Considering cell count alone, 34 (20%) of 171 compounds resulted in cytotoxicity at 10 μM in human microglial cells, while 67 (39%) resulted in cytotoxicity at 10 μM in mouse microglial cells. Except for three compounds, all compounds that were toxic to human microglial cells were also toxic to mouse microglial cells. This included several compounds targeting kinases (ALK, CDKs, MARK4, and SYK), FOXO1, and VCP. Gratifyingly, of the 11 positive control compounds, only a compound with observed red fluorescence (UNC10303097A/ TRx0237 (LMTX) mesylate) that likely interfered with the assay manifested toxicity in both microglial cell lines. UNC10126573A/ masitinib (AB1010) displayed cytotoxicity only in mouse microglia. The remainder of compounds optimized for clinical use for AD did not exhibit cytotoxicity, which supports that optimization of the chemical leads in the AD Informer Set can reduce toxicities resulting from off-target liabilities.

For phagocytosis, <70% control was considered inhibition, while >150% control was considered stimulation. It’s important to note that stressed cells, such as those that are dying, are prone to show an increase in phagocytosis. When considering the phagocytosis results in human microglial cells, many more compounds were inhibitory (71) than stimulatory (7). Most compounds that demonstrated cytotoxicity (cell count <70% control) were also inhibitors of microglial phagocytosis at 10 μM, while a few fell into the stimulator category. When considering the phagocytosis results in mouse microglial cells, the opposite trend was observed: many more compounds were stimulatory (70) than inhibitory (33). Efforts are underway to understand these differences in the response of human (HMC3) and murine (BV2) to treatment in order to pick the best cell line to use in assays moving forward. As was observed in the human cells, most compounds that demonstrated cytotoxicity (cell count <70% control) were also inhibitors of microglial phagocytosis at 10 μM and a few were stimulators. Only one compound (UNC10302867A/ MRT67307) was a stimulator of phagocytosis in both human and mouse microglia, while many compounds were inhibitors of phagocytosis in both human and mouse microglia. When considering the published literature that connects gene targets with phagocytosis in the context of AD, DOCK1 (UNC8367A/ TBOPP) has been reported to play a role in phagocytosis and was found to be stimulatory in human microglia at 10 μM in our assay as well [51]. MerTK promotes phagocytosis and thus MerTK inhibition is proposed to lead to inhibition of phagocytosis [52]. In accordance with this hypothesis, MERTK inhibitor UNC2025C (same as UNC2025) [53] inhibited mouse and human phagocytosis in our assay at 10 μM.

In the context of AD therapy, we are looking for compounds that can stimulate microglial phagocytosis with low or no associated cellular toxicity. With these parameters in mind and based on the single-concentration data, several compounds were selected for dose-response follow-up. These compounds were initially dosed up to 20 μM for 24h. It was clear that trends were emerging at the highest doses, so then the dose range was expanded up to 40 μM for 24h for a second round of testing. Finally, since it takes time for some targets to respond to treatment and for changes in gene expression to manifest, we next dosed up to 40 μM for 48h.

UNC10302865A/ sapitinib (AZD8931), UNC10240506B/ donepezil hydrochloride (E2020), and UNC10100724A/ ketoconazole were selected for the first round of dose-response follow-up because they stimulated phagocytosis in one of the cell lines tested with no or low associated cytotoxicity in the single concentration testing (Table 1). When we compare the data from Table 1 and Figure 1, panel A for UNC10302865A/ sapitinib (AZD8931), we see that the stimulation of microglial phagocytosis in HMC3 cells and inhibition of phagocytosis in BV2 cells predicted by the single-concentration data (Table 1) was observed in dose-response when tested up to 20 μM (top graphs) and was even more striking when the concentration range was expanded up to 40 μM (bottom graphs) following 24h treatment. Some toxicity was observed at the highest concentrations in BV2 cells. For UNC10240506B/ donepezil hydrochloride (E2020), the single concentration data predicted stimulation of phagocytosis in both cell lines (Table 1). Robust stimulation was observed in both cell lines when tested in dose-response (Figure 1, panel B), with toxicity limited only to the 40 μM dose in BV2 cells. For both UNC10302865A/ sapitinib (AZD8931) and UNC10240506B/ donepezil hydrochloride (E2020), there is clearly a dosing window between observed toxicity and a phenotypic response. Also, in both cases, the single-point data was predictive of the dose-response data.

**Table 1.** Summary of 10 μM single-concentration data, normalized to control (DMSO) treated cells, for compounds selected for dose-response follow-up.

|  | Gene target | Cell number |  | Nuclear size |  | DNA intensity |  | Phagocytosis |  |
| --- | --- | --- | --- | --- | --- | --- | --- | --- | --- |
|  |  | HMC3 | BV2 | HMC3 | BV2 | HMC3 | BV2 | HMC3 | BV2 |
| **UNC10302865A** | ERBB3 (HER3) | 132.1 | 102.4 | 105.8 | 110.1 | 90.8 | 101.0 | 152.7 | 68.8 |
| **UNC10240506B** | ACHE | 130.2 | 126.0 | 100.5 | 103.1 | 98.0 | 104.5 | 123.4 | 255.2 |
| **UNC10100724A** | CYP3A43 | 125.5 | 129.7 | 96.3 | 103.8 | 100.1 | 104.7 | 144.4 | 280.0 |
| **UNC10244898A** | ALK | 127.2 | 117.2 | 110.3 | 102.7 | 89.2 | 110.1 | 37.8 | 200.3 |

**Figure 1.**
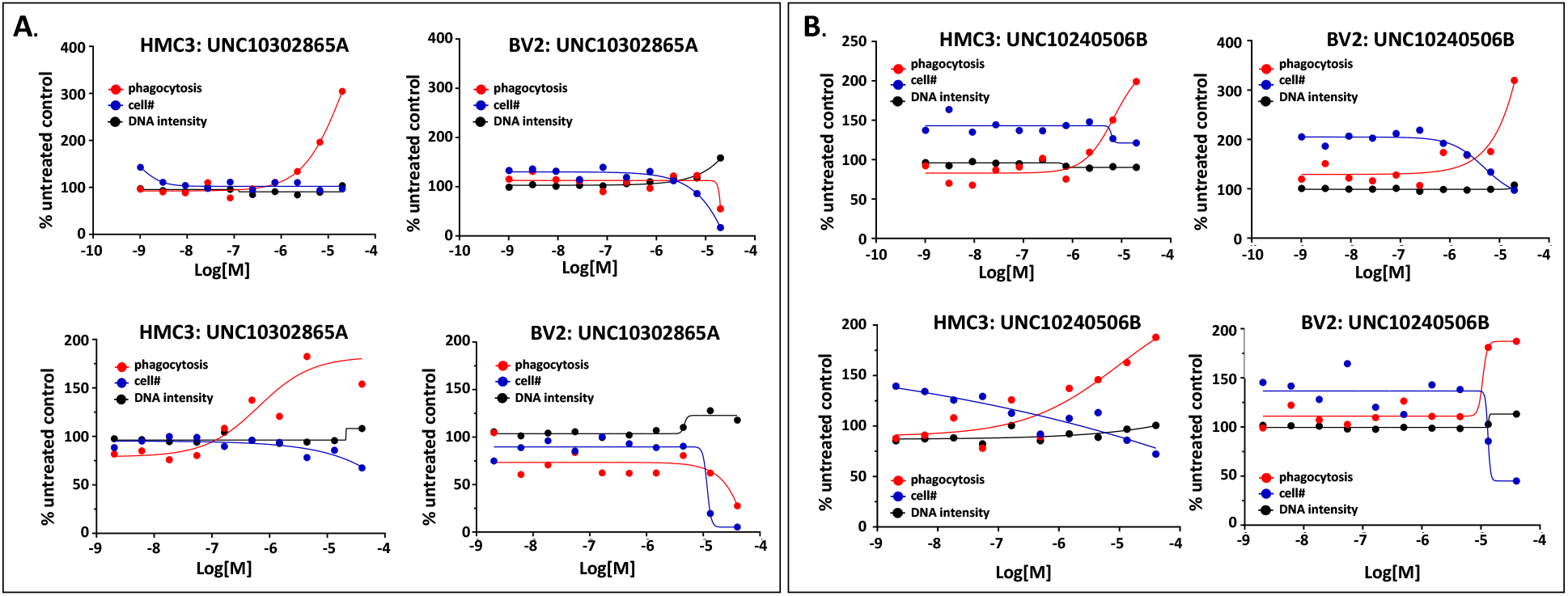
Phagocytosis assay 24h dose-response follow-up for UNC10302865A and UNC10240506B in HMC3 and BV2 cells.

Like the compounds in Figure 1, UNC10100724A/ ketoconazole was dosed up to 20 μM (Figure 2, panel A, top graphs) and up to 40 μM (Figure 2, panel A, bottom graphs) for 24h. The same compound was also dosed up to 40 μM for 48h (Figure 2, panel B). The single-concentration data (Table 1) predicted stimulation of phagocytosis in both cell lines. This stimulation of phagocytosis was observed when cells were treated for either 24h or 48h with UNC10100724A in dose-response and without notable toxicity. Finally, UNC10244898A/ lorlatinib was introduced in dose-response up to 40 μM for 48h (Figure 2, panel C). As shown in Table 1, UNC10244898A was predicted by single-concentration data to inhibit phagocytosis in HMC3 cells and stimulate it in BV2 cells. Some toxicity was observed with UNC10244898A at the highest doses, but inhibition/stimulation of phagocytosis occurs at a much lower concentration and provides a dosing window to elicit these responses without associated toxicity. Mechanisms driving the observed differential phagocytic responses to UNC10244898A in HMC3 and BV2 cells are unknown and warrant further study. The 48h dose-response data once again reinforces that the single-concentration data was predictive of the phagocytic phenotypic response to be observed when a compound is dosed over a broad concentration range. This comparison of single and dose-response data in our phagocytosis assay gives us confidence that the single-concentration data can be used to reliably predict a phagocytic phenotype using our assay system.

**Figure 2.**
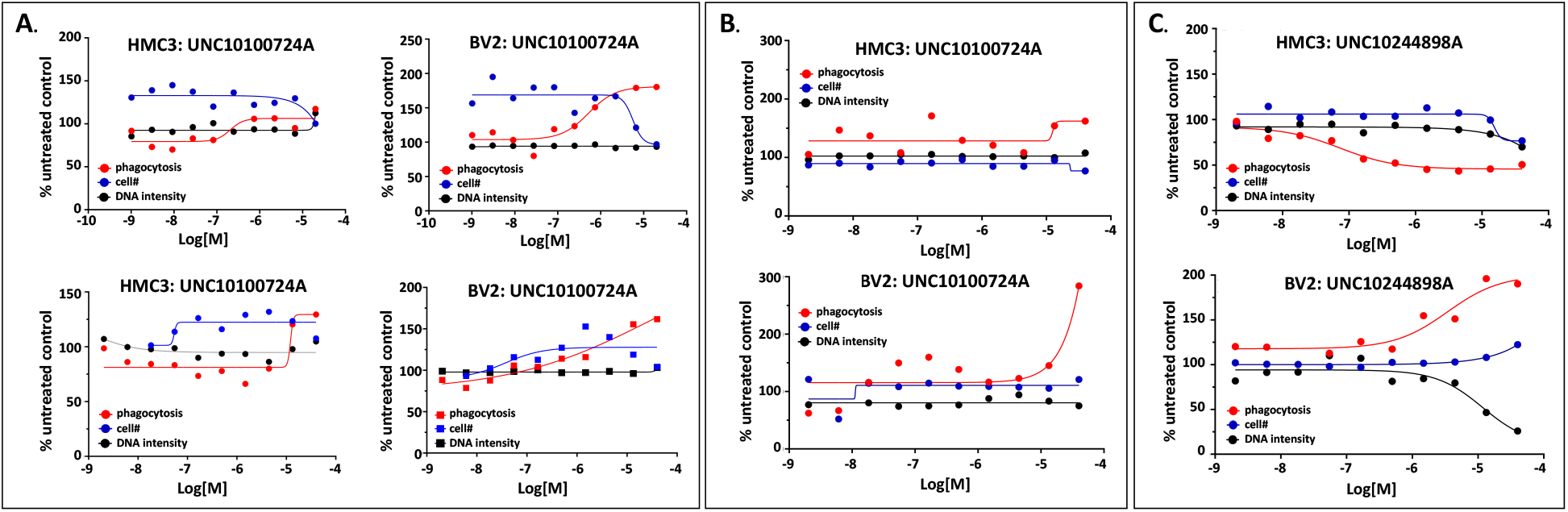
Phagocytosis assay 24h or 48h dose-response follow-up for UNC10100724A and UNC10244898A in HMC3 and BV2 cells.

G protein-coupled receptors (GPCRs) are highly expressed, essential receptors in the brain involved in processes such as neuronal communication, neurogenesis, movement, and cognition [54-56]. Given their abundance and importance, GPCRs could present potential off-target liabilities for compounds of AD targets in the brain. Analysis of the GPCR panel screening results yielded many interesting findings (Figure 3). UNC10302682A/ P505-15 (PRT06207 HCl) and UNC2025C possessed affinity for the most receptors in primary assays and were thus analyzed in the most secondary assays. Confirmed affinity (via secondary assays) was only observed for a portion of the receptors identified in the primary assays. UNC10302682A/ P505-15 (PRT06207 HCl) possessed modest affinity for the Sigma2 (3.0 μM), Alpha1A (5.1 μM), H1 (1.2 μM), 5-HT2C (7.2 μM), 5-HT2A (7.2 μM), 5-HT1D (3.3 μM), and 5-HT3 (1.9 μM) receptors, and the norepinephrine and dopamine transporters (1.0 and 2.1 μM, respectively). In contrast, UNC2025C possessed affinity for the Sigma2 (0.2 μM), Alpha1D (5.0 μM), H4 (4.0 μM) and 5-HT2A (0.4 μM) receptors, and the norepinephrine, serotonin and dopamine transporters (1.4, 0.9 and 9.9 μM, respectively). While these compounds demonstrated affinity for several GPCRs, it is worth noting that their on-target activity provides a large window at which to dose without engaging these peripheral receptors. The Ki of UNC2025C for MERTK, for example, is ∼200 pM while the Ki values observed in these secondary GPCR assays were in the micromolar range [53]. Since UNC10302682A/ P505-15 (PRT06207 HCl) and UNC10302678A/ entospletinib (GS-9973) are both inhibitors of SYK, the GPCR affinity observed for UNC10302682A/ P505-15 (PRT06207 HCl) does not seem tied to SYK inhibition but rather to off-target protein binding interactions. Similarly, another published SYK inhibitor (BI 1002494) also lacked this activity when profiled against many of the same GPCRs [57]. Interestingly, UNC10302682A/ P505-15 (PRT06207 HCl) and UNC2025C are both kinase inhibitors. Their structures, however, are not very similar. With a few exceptions, compounds targeting EPHX2, CAPN2, and MDK did not exhibit high affinity in secondary binding assays.

**Figure 3.**
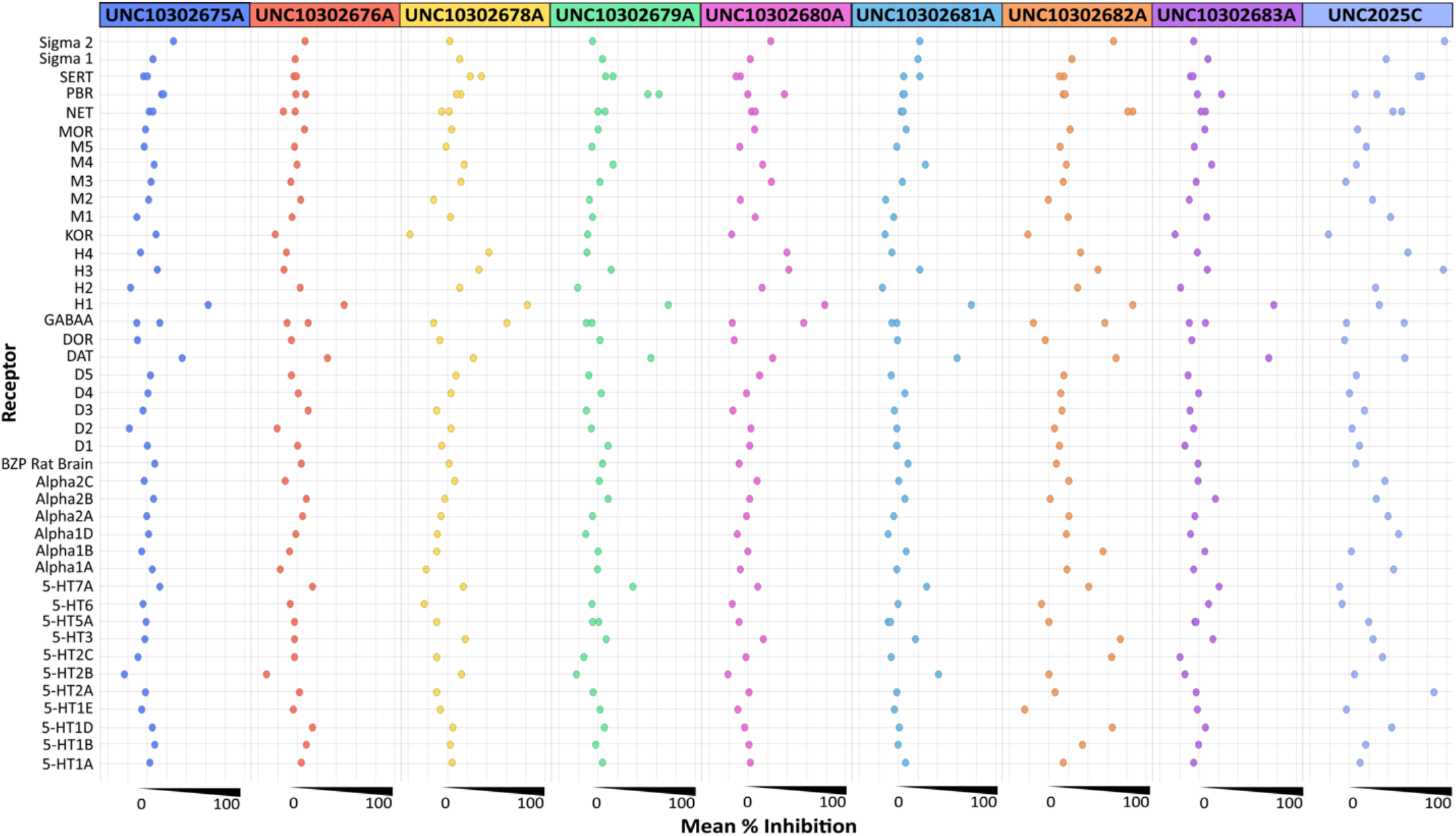
Visualization of GPCR primary binding data for selected AD Informer Set compounds.

Our initial iPSC-based neuronal studies provide an example of testing AD Informer Set compounds in primary cell assays. These preliminary screening results demonstrate the utility of the set and offer the context upon which to carry out subsequent mechanistic studies. Aβ40 and Aβ42 are secreted in response to the three major cleavages (γ, ε, and ζ) within the transmembrane domain of APP [58]. While Aβ40 is more prevalent, the relative ratios of Aβ42:40 can be calculated to determine if there is a specific effect of a compound on the γ-secretase. For example, some familial AD mutations in presenilin-1 cause a change in γ-secretase cleavage such that Aβ42 peptides are increased and Aβ40 peptides are decreased [59]. However, for this study, we did not observe a change in Aβ42:40 ratios upon treatment (Figure 4). In some cases, we detected fluctuations in individual Aβ peptide levels, for example with UNC10302679A, UNC10302681A, UNC10302683A, and UNC10302680A at 24hrs (Figure 4C and E) however by 48 hours of treatment most of these had resolved (Figure 4D and F).

**Figure 4.**
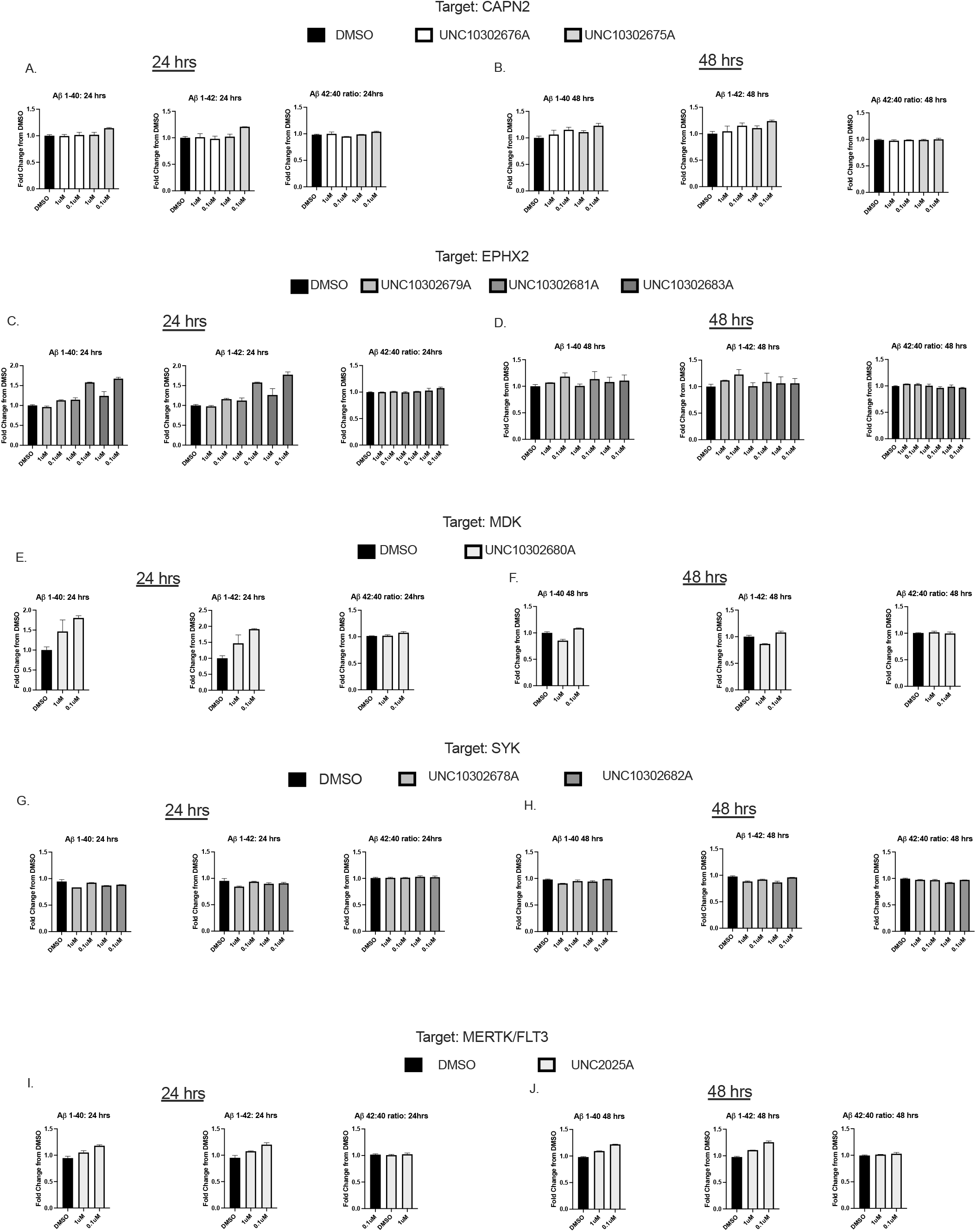
Aβ peptide secretion assay results for (A-B): UNC10302676A and UNC10302675A; (C-D):UNC10302679A, UNC10302681A, UNC10302683A; (E-F): UNC10302680A; (G-H): UNC10302678A, UNC10302682A; (I-J): UNC2025C. Aβ40 is in the left panel, Aβ42 in the middle panels and the Aβ42:48 ratio in the bottom panel. Indicated are 24hr (A,C,E,G,I) and 48hr (B,D,F,H,J) treatment times.

The Aβ secretion results (Figure 4) demonstrated that compounds targeting CAPN2, EPHX2, MDK, SYK, or MERTK/FLT3 do not have a strong effect on Aβ40, Aβ42 or the resultant ratio of the two peptides. This finding was true at both concentrations examined and at both time points as well. In a previous report, UNC10302675A/ MDL 28170 did not inhibit the secretion of Aβ40 and Aβ42 in N2a cells stably expressing wild-type presenilin 1 and myc-tagged Swedish mutant APP when dosed at 30 μM for 24h. Levels of secreted Aβ40 and Aβ42 increased dramatically between 24 and 48h at this concentration. At concentrations <5 μM, treatment for 48h did not result in a response in secreted Aβ40 or Aβ42, supporting our results [58]. EPHX2 inhibitors have been reported to prevent the cytotoxicity induced by Aβ peptides, but there is no report on their effect on modulating secreted Aβ levels [60]. The same is true for MDK, which binds directly to Aβ40 to inhibit its cytotoxicity *in vitro* [61]. SYK inhibitor BAY61-3606 (2mg/kg) stimulated transport of Aβ40 and Aβ42 in transgenic PS1/APPsw mice, resulting in a significant reduction in the detectable levels of both peptides [62].

The tau phosphorylation assay results were equally interesting. Excessive phosphorylation of tau is a hallmark for AD and thus agents that reduce tau phosphorylation are sought [63]. A decrease in the ratio of pTau:tTau suggests a decrease in tau phosphorylation. In this study, when we calculated the pTau:tTau ratios, we observed that several compounds reduced the pTau:tTau ratio in the neurons (Figure 5, indicated by black arrows). In particular, a reduced ratio was observed for UNC10302681A/ TPPU, UNC10302683A/ AR9281, and UNC10302680A/ iMDK when dosed at 1 μM for 24h. This decrease was not observed at the lower concentration after 24h or at 48h for either concentration. UNC10302681A/ TPPU and UNC10302683A/ AR9281 target EPHX2, while UNC10302680A/ iMDK is an MDK inhibitor. In support of our results, treatment of AD model mice with UNC10302681A/ TPPU (5 mg/kg/day) resulted in a reduction of tau hyperphosphorylation species (Ser396 and Ser404) [64]. Ebselen, an irreversible inhibitor of EPHX2, has also been shown to decrease tau phosphorylation in a triple transgenic AD mouse model [63, 65]. It has been suggested that oxidative stress can promote tau hyperphosphorylation and resultant aggregation and that inhibition of EPHX2 is anti-inflammatory [64]. Less has been published establishing a connection between MDK and tau phosphorylation. No effect on the pTau:tTau ratio was observed at either concentration or time point for compounds targeting CAPN2, SYK, or MERTK/FLT3.

We further examined the total levels of each peptide (Figure 6).Interestingly, at the higher 1 μM dose, EPHX2 inhibitors UNC10302681/ TPPU and UNC10302683A/ AR9281 decreased both pTau and tTau but the decrease in pTau was more pronounced, resulting in an observed shift in the ratio (Figure 6C, indicated by black arrows). UNC10302680A/ iMDK also showed a larger decrease in pTau (Figure 6E, indicated by an arrowhead). While this was observed at 24h, the effect was not as pronounced at 48h, which leads us to suggest that this is not due to toxicity of the compounds. With SYK inhibitors, we observed a decrease in total levels of pTau and tTau at 48 hours with both doses (Figure 6H, indicated by dashed arrows). This is in line with previous reports showing that SYK inhibition leads to decreased tau expression and a reduction in phospho-tau [66]. It has been suggested that inhibition of SYK with the inhibitor BAY61-3606 indirectly reduces tau phosphorylation via GSK3β and/or PI3K inhibition, but this result was only observed at higher concentrations than we tested (10 μM) [62]. In our treatments, both tTau and pTau were decreased to the same extent, therefore not changing the pTau:tTau ratio. While FLT3 has been implicated as a kinase that phosphorylates tau *in vitro*, there is no report of FLT3 inhibitors impacting tau phosphorylation [67]. Similar to the reported mechanism for SYK, an indirect link has been proposed that links CAPN2 activation and tau phosphorylation in AD via CDK5 activation [68, 69].

**Figure 5.**
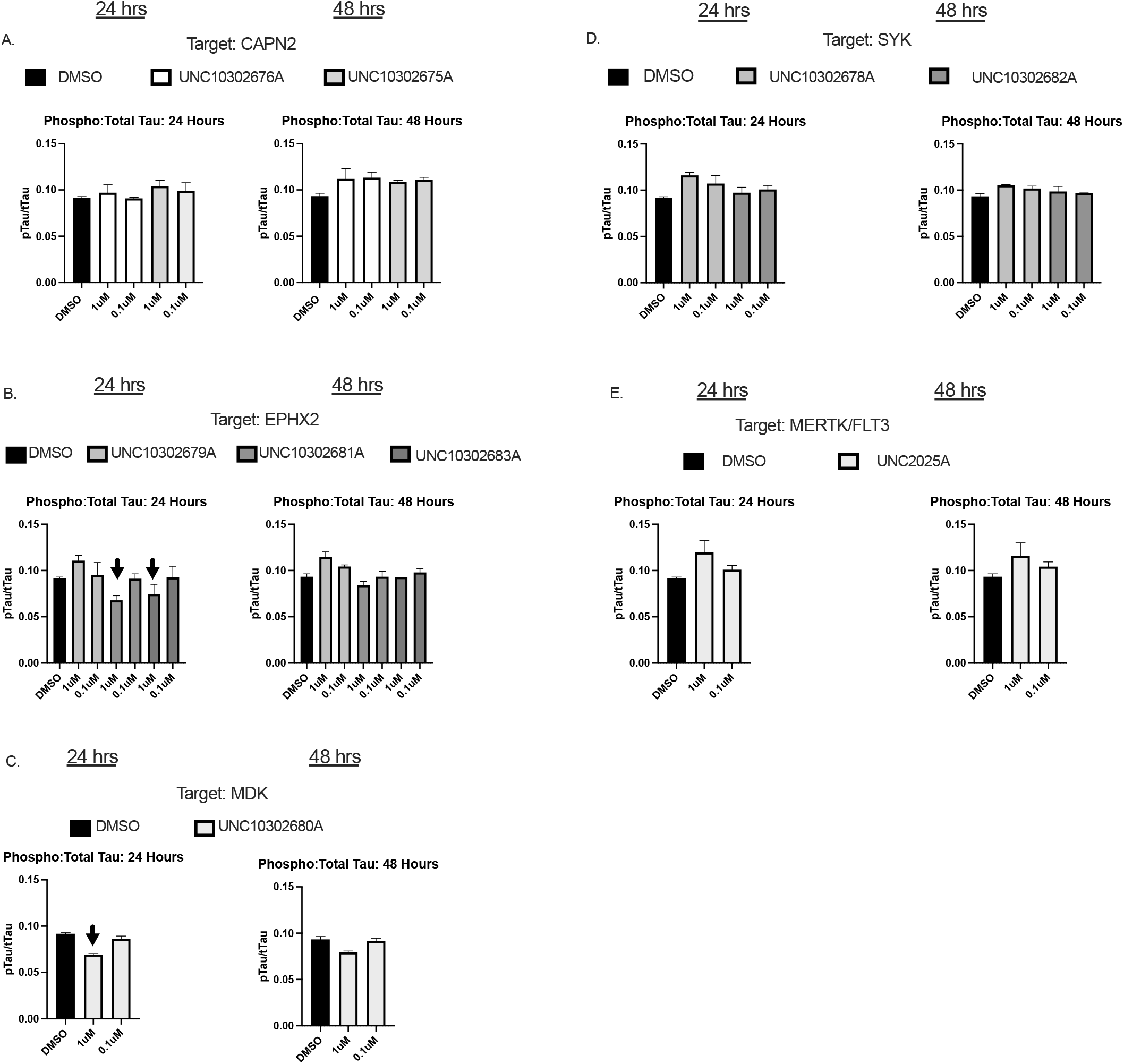
Phospho:total tau ratio results for (A): UNC10302676A and UNC10302675A; (B):UNC10302679A, UNC10302681A, UNC10302683A; (C): UNC10302680A; (D): UNC10302678A, UNC10302682A; (E): UNC2025C. UNC10302681A, UNC10302683A and UNC10302680A lowered the pTau:tTau ratio at the 1 μM dose after 24 hours of treatment (panels B and C, indicated by black arrows). 24hr and 48hr timepoints are indicated.

**Figure 6.**
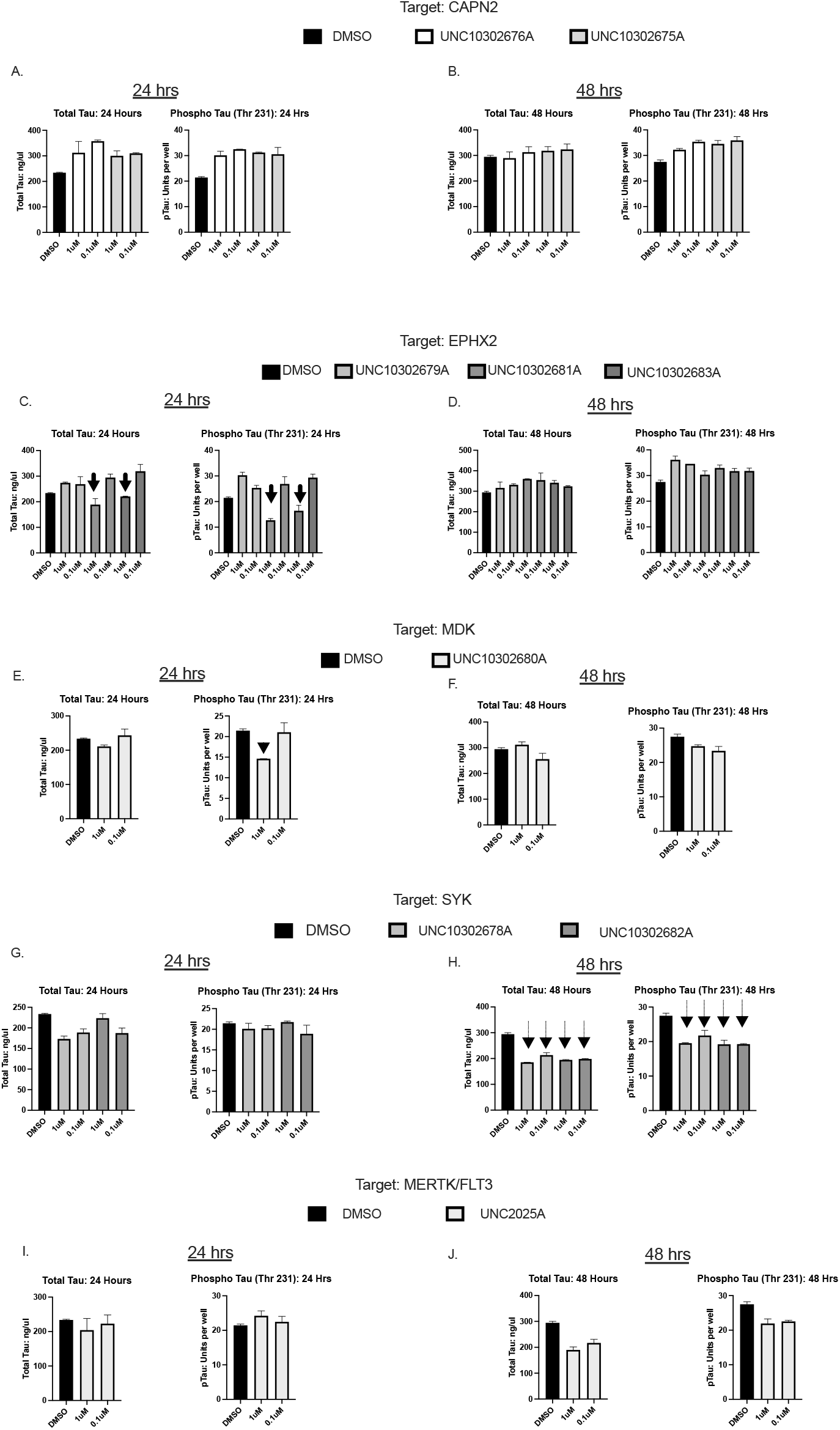
Results for individual levels of phospho and total tau peptides for (A-B): UNC10302676A and UNC10302675A; (C-D):UNC10302679A, UNC10302681A, UNC10302683A; (E-F): UNC10302680A; (G-H): UNC10302678A, UNC10302682A; (I-J): UNC2025C. UNC10302681A and UNC10302683A lowered both pTau and tTau with a larger effect on the pTau levels (panel C, indicated by black arrows). UNC10302680A reduced the pTau levels with little effect on tTau (panel E, black arrowhead) and UNC10302682A reduced pTau and tTau peptides to the same extent (panel H, dashed arrows). 24hr and 48hr timepoints are indicated.

Our hiPSC-derived neuronal model provides a platform where these compounds can be screened for effects on AD-related peptides in a biologically relevant system. It is our goal that these preliminary results will provide baseline information for other investigators to perform more detailed analyses with these compounds. To further probe our findings, we will use the RNA harvested from this experiment and use qRT-PCR to examine specific target genes.

The mouse PK experiments, summarized in Table 2, taught us more about specific compounds in the AD Informer Set. Firstly, UNC10302681A/ TPPU demonstrated excellent PK with a long half-life (T_1/2_ = 13.8 h) and low clearance (1.3 mL/min/kg). In contrast, UNC10302683A/ AR9281 revealed poor PK with a short half-life (T_1/2_ = 0.24 h) and high clearance (93 mL/min/kg), and we could not check brain permeability. Based on these findings, UNC10302681A/ TPPU was selected for follow-up studies aimed at determining its brain permeability while UNC10302683A/ AR9281 was excluded. The remaining four compounds sent for snapshot PK exhibited half-life and clearance values in between these two. Since UNC10302678A/ entospletinib (GS-9973) and UNC10302680A/ iMDK required formulation in NMP/PEG-400 (10:90) to execute the snapshot PK studies and this formulation cannot be used *in vivo*, studies aimed at determining their brain penetration were not pursued. Despite its solubility issues that required this formulation, which were also confirmed in kinetic solubility experiments, UNC10302678A/ entospletinib (GS-9973) is in Phase 2 clinical trials for hematological malignancies. UNC10302680A/ iMDK also demonstrated poor solubility in the kinetic solubility experiment but has been dosed in mice [70]. UNC10302681A/ TPPU, has been dosed systemically in many preclinical animal models with chronic inflammatory conditions, while UNC10302683A/ AR9281, despite its short half-life, is in Phase 2 clinical trials for patients with hypertension and impaired glucose tolerance [71-73]. UNC10302679A/ GSK2256294A and UNC10302682A/ P505-15 (PRT06207 HCl) were also advanced as far as Phase 2 clinical trials for insulin resistance and subarachnoid hemorrhage (UNC10302679A), and rheumatoid arthritis (UNC10302682A).

**Table 2.** Summary of published and experimentally determined IV mouse PK values.

| Compound ID | Dose (mg/kg) | $T_{1/2}$ (h) | $C_{max}$ (μM) | $AUC_{last}$ (h*μM) | CL (mL/min/kg) | $V_{ss}$ (L/Kg) | Brain: Plasma at 1h (10 mg/kg) |
| --- | --- | --- | --- | --- | --- | --- | --- |
| UNC10302676A* | 2.0 | 6 | - | 35.6 | 7.8 | 1.0 | 0.015 |
| **UNC10302675A\*** | 3.0 | 0.74 | 2345 | 1184 | 41.8 | 1.23 | 0.10 |
| **UNC2025C\*** | 3.0 | 3.8 | 4.36 | 9.78 | 9.22 | 2.33 | 1.96 |
| **UNC10302679A** | 3.0 | 0.64 | 4011 | 1892 | 26.3 | 0.62 | 0.018 |
| **UNC10302681A** | 3.0 | 13.8 | 2144 | 7149 | 1.31 | 1.55 | 0.46 |
| **UNC10302683A** | 3.0 | 0.24 | 1832 | 537 | 93.1 | 0.47 | ND |
| **UNC10302680A** | 3.0 | 1.16 | 1569 | 1240 | 37.8 | 2.61 | ND |
| **UNC10302682A** | 3.0 | 1.64 | 595 | 831 | 49.7 | 6.26 | 0.08 |
| **UNC10302678A** | 3.0 | 0.90 | 926 | 1674 | 28.5 | 2.34 | ND |

Abbreviations: AUC<sub>max</sub>, maximal area under the curve (drug concentration as a function of time); Brain:Plasma, brain to plasma ratio; CL, drug clearance; C<sub>max</sub>, highest concentration of drug measured; ID, identifier; T<sub>1/2</sub>, half-life; V<sub>ss</sub>, steady state volume of distribution.
Notes: \*Published PK values and experimentally determined Brain:Plasma ratios.

Studies aimed at experimentally determining the brain exposure at 1h for compounds with published and acceptable PK highlighted UNC2025C as the most promising. UNC2025C demonstrated a brain:plasma ratio of 1.96 [74]. All other compounds demonstrated brain:plasma ratios <1 and ranged from 0.015 for UNC10302676A/ ABT-957 (Alicapistat) to 0.46 for UNC10302681A/ TPPU. For the compounds targeting EPHX2 (UNC10302679A/ GSK2256294A, UNC10302681A/ TPPU, and UNC10302683A/ AR9281), no BBB data was found for UNC10302679A, but UNC10302681A and UNC10302683A were reported as brain penetrant [64, 71, 72, 75]. The brain-to-plasma ratio of UNC10302681A/ TPPU when dosed orally in mice at 3 mg/kg was reported as 0.18 [60, 76]. Baboon PET studies were performed with blocking doses of UNC10302683A/ AR9281 and levels in the brain versus plasma were quantified as well as specific regions of the brains imaged [75]. No reported BBB data could be found for the compounds targeting SYK (UNC10302682A/ P505-15 (PRT06207 HCl)) or CAPN2 (UNC10302676A/ ABT-957 (Alicapistat) and UNC10302675A/ MDL 28170). With these experimental values, we attempted to determine whether any of the methods used to predict BBB penetration was more accurate and could be relied upon in compound design. Three different predictive methodologies were compared: CNS MPO (Central Nervous System MultiParameter Optimization), the StarDrop software program [28], and the ADMET Predictor software program [31]. CNS MPO, developed and utilized by scientists at Pfizer, is calculated based upon 6 parameters: (1) lipophilicity, calculated partition coefficient (ClogP); (2) calculated distribution coefficient at pH=7.4 (ClogD); (3) molecular weight (MW); (4) topological polar surface area (TPSA); (5) number of hydrogen bond donors (HBD); and (6) most basic center (pKa). A CNS MPO score ≥4 suggests that a compound will be BBB permeable [77].

StarDrop is software for small molecule design and optimization that uses compound physicochemical properties to predict the concentration of compound in the brain versus in the blood as an expectation of BBB penetration [28]. A BBB log([brain]:[blood]) >-0.5 is predicted to indicate BBB penetrance. ADMET Predictor is a machine learning software tool that predicts over 175 properties, divided into eight different modules, based simply on the 2D structure of the molecule [31]. The Physicochemical and Biopharmaceutical module includes a predictor of whether a compound will penetrate the BBB as well as a calculation of the logarithm of the brain/blood partition coefficient (RMSE/MAE = 0.37/0.28). The literature reports slightly different logBB cutoffs to classify compounds as either BBB penetrant or not. These values range from -1.00 to 0.63, with many opting to report the cutoff criteria to be BBB penetrant as logBB = 0.00 (i.e., compounds with logBB ≥ 0.00 are predicted to be BBB penetrant) [78-80].

As seen in Table 3, the calculated predictive methods did not always agree with one another, only UNC10302681A/ TPPU and UNC2025C were suggested to be brain penetrant by two of the three methods, and the rank order of brain:plasma ratio was not accurately predicted by any of them. At least one of the predictive tools indicated that UNC2025C, UNC10302675A/ MDL 28170, and UNC10302681A/ TPPU (values highlighted in green in Table 3) would cross the BBB and these three compounds showed the highest brain:plasma ratios when determined experimentally. We suggest that consideration of calculated scores from multiple sources is the optimal practice moving forward, with the understanding that if two predict brain permeability then chances of experimental verification are increased. This first iteration of the AD Informer Set is available immediately to the research community. We anticipate supplementing the set and releasing additional compounds with associated data as part of future iterations. We encourage interested individuals to use the set, publish their results, and we will do our best to aid in data deconvolution, analysis and further annotation of this resource.

**Table 3.** Experimental versus calculated BBB penetration values.

| Compound ID | Brain:Plasma at 1h<br>(10 mg/kg) | MPO score | StarDrop:<br>BBB $\log([brain]/[blood])$<br>value | ADMET Predictor:<br>LogBB value |
| --- | --- | --- | --- | --- |
| UNC2025C | 1.96 | 2.7 | -0.1211 | 0.25 |
| UNC10302676A | 0.015 | 3.8 | -1.006 | -0.438 |
| UNC10302675A | 0.10 | 3.3 | -0.2819 | -0.408 |
| UNC10302679A | 0.018 | 2.6 | -0.6425 | -0.386 |
| UNC10302681 | 0.46 | 5.3 | -0.4012 | -0.261 |
| UNC10302682 | 0.08 | 2.5 | -0.7396 | -0.237 |

Abbreviations: BBB, blood-brain barrier; Brain:Plasma, brain to plasma ratio; ID, identifier; LogBB and $\log([brain]/[blood])$ , logarithmic ratio between the concentration of a compound in brain and blood; MPO, MultiParameter Optimization score.

## Supporting information

GPCR data

Microglia data

Plate map

Annotation data

## ACKNOWLEDGMENTS

The authors thank the collaborative TREAT-AD centers headed by Allan Levey and Alan Palkowitz, the TREAT-AD advisory board members, as well as Scientific Officers within NIA for lending their expertise to this study: Lara Mangravite, Haian Fu, Opher Gileadi, Aled Edwards, Frank Longo, and Gregory Carter. Primary and secondary GPCR binding assays were generously provided by the National Institute of Mental Health’s Psychoactive Drug Screening Program, Contract # HHSN-271-2018-00023-C (NIMH PDSP). The NIMH PDSP is directed by Bryan Roth at the University of North Carolina at Chapel Hill and Project Officer Jamie Driscoll at NIMH in Bethesda, MD.

## SUPPORTING INFORMATION

Additional supporting information may be found online in the Supporting Information section at the end of the article.

